# Phenotypically Complex Living Materials Containing Engineered Cyanobacteria

**DOI:** 10.1101/2023.01.26.525792

**Authors:** Debika Datta, Elliot L. Weiss, Daniel Wangpraseurt, Erica Hild, Shaochen Chen, James W. Golden, Susan S. Golden, Jonathan K. Pokorski

## Abstract

A cyanobacterial photosynthetic biocomposite material was fabricated using 3D-printing and bioengineered to produce multiple functional outputs in response to an external chemical stimulus. Our investigations show the advantages of utilizing additive manufacturing techniques in controlling the design and shape of the fabricated materials, which proved to be important for the support and growth of obligate phototrophic microorganisms within the material. As an initial proof-of-concept, a synthetic theophylline-responsive riboswitch in *Synechococcus elongatus* PCC 7942 was used for regulating the expression of a yellow fluorescent protein (YFP) reporter. Upon induction with theophylline, the encapsulated cells produced YFP within the hydrogel matrix. Subsequently, a strain of *S. elongatus* was engineered to produce an oxidative enzyme that is useful for bioremediation, laccase, expressed either constitutively or under the control of the riboswitch. The responsive biomaterial can decolorize a common textile dye pollutant, indigo carmine, potentially serving as a useful tool in environmental bioremediation. Finally, cells were engineered to have the capacity for inducible cell death to eliminate their presence once their activity is no longer required, which is an important function for biocontainment and minimizing unintended environmental impact. By integrating genetically engineered stimuli-responsive cyanobacteria in patterned volumetric 3D-printed designs, we demonstrate the potential of programmable photosynthetic biocomposite materials capable of producing functional outputs including, but not limited to, bioremediation.

## Introduction

Synthetic stimuli-responsive polymeric materials have been fabricated to sense and respond to different environmental conditions such as chemicals, pH, light, and temperature. Such polymers have been utilized for applications including therapeutics^1^, drug delivery^2^, biomedical devices^3^, biosensors^4,5^, electronics^6,7^, and soft robotics^8^. However, in most cases, the ability of these materials to respond with high specificity and produce the desired output is limited by the nature of the available stimuli and is restricted by the specifics of a chemically driven response. The development of purely synthetic materials is therefore limited by the coupling of available external cues as well as the extent and nature of the material’s response.

Unlike traditional materials systems, biological systems innately respond and adapt to their surrounding environment and can generate bioproducts with a wide range of functions. The field of synthetic biology aims to exploit these natural systems by engineering them to perform high-value user-defined functions for the production of biofuels, proteins, fertilizers, bioplastics, and catalysts^9–13^. Recent progress in synthetic biology enables the design and manipulation of cellular signaling pathways for regulating gene expression for the constitutive or induced (in response to an environmental stimulus) production of a desired biosynthetic output.

The emerging field of engineered living materials (ELMs) aims to design programmable materials^14–21^. Compared to synthetic stimuli-responsive materials, ELMs use genetically engineered biological components, integrated into a composite material, to produce functional outputs in response to environmental cues. Recent ELMs have utilized a diversity of microorganisms including bacteria, yeast, fungi, and algae. Some notable examples include the fabrication of skin patches for wound healing^22^, sweat-responsive biohybrid fabrics^23^, a biodegradable aqua plastic^24^, the incorporation of chloroplasts for self-repairing hydrogels^25^, and photosynthetic O_2_ generators for enhancing mammalian cell viability^26^. While the potential of ELMs is vast, there are significant challenges in their development including the necessity for genetically stable engineered strains, the formulation of materials to support the growth and viability of engineered cells over long durations of time, and programming of cells to be stimuli-responsive at the material interface. Thus, an appropriate pairing of biological components and polymeric materials is critical to developing any stimuli-responsive ELM.

With the aim of developing innovative sustainable materials and overcoming the limitations of traditional stimuli-responsive materials, we chose to incorporate photosynthetic cyanobacterial cells within a polymer composite material. Cyanobacteria are appealing candidates for inclusion in ELMs due to their viability in low-cost media, with the added benefit that the carbon source for these phototrophs, CO_2_, is freely available. Moreover, the abundance of genetic engineering tools available for several cyanobacterial species allows for modular systems of plasmid design and gene expression controlled by external cues. Engineered plasmids can be self-replicating or designed for integration of heterologous genes into the chromosome.

Over the last decade, a number of tools have been developed and used for genetic modification in cyanobacteria and applied for heterologous expression of high-value products and regulatory circuits^27–29^. Notably, the development of various cyanobacteria-compatible promoters, ribosomal binding sites (RBSs), reporters, modular vectors, and markerless selection systems have contributed greatly to the evolution of cyanobacterial synthetic biology^30,31^. Of relevance to this study is the use of a riboswitch as a tool for regulating gene expression. Riboswitches are aptameric sequences in the mRNA that regulate gene expression in response to ligand binding and can be used to alter rates of protein production. Previous studies have demonstrated the utility of the theophylline-responsive riboswitch-F in the cyanobacterium *Synechococcus elongatus* PCC 7942 (*S. elongatus*), which, when supplemented with the small molecule theophylline, undergoes a conformational change allowing the mRNA to bind to an RBS, resulting in the translation of a target protein^32^. In the stimuli-responsive material presented in this study, the translation of proteins from heterologous genes of interest is regulated in the cyanobacteria-polymer composite using a chemically responsive synthetic riboswitch.

Bioremediation is an area where ELMs containing genetically modified organisms are likely to provide substantial benefit^33,34^. Previous work has demonstrated that cyanobacteria are capable of producing both native and recombinant enzymes with relevance to bioremediation^35–37^. In this work, we aimed to develop stimuli-responsive ELMs that decontaminate chemical pollutants by regulating the production of an enzyme capable of degradation. Furthermore, we engineered cyanobacteria to have a ‘kill switch’ and show the induction of lytic cell death upon riboswitch activation, limiting biofouling in the case of cells leaching from the ELM. The work presented herein is a sustainable alternative to develop materials that respond to chemical stimuli with phenotypically complex outputs beyond that which could be achieved in a fully synthetic system.

## Results and discussion

### Fabrication of ELM with *S. elongatus* and alginate hydrogel

Several ELMs were fabricated that contain engineered strains of *S. elongatus* expressing heterologous proteins, and the material properties, cell viability, and biological activity of the composite materials were investigated. ELMs were constructed to contain *S. elongatus* strains engineered to express a variety of proteins under the regulation of a theophylline-responsive riboswitch to demonstrate stimuli-responsive photosynthetic biomaterials. **Figure 1** illustrates a schematic overview of the fabrication of these ELMs that combined genetically-engineered cyanobacterial cells with an alginate polymer to create a 3D-printed hydrogel composite material; the gene of interest in the cells is activated by the presence of an inducer molecule (input) producing a specific protein of interest (output).

**Figure 1.**
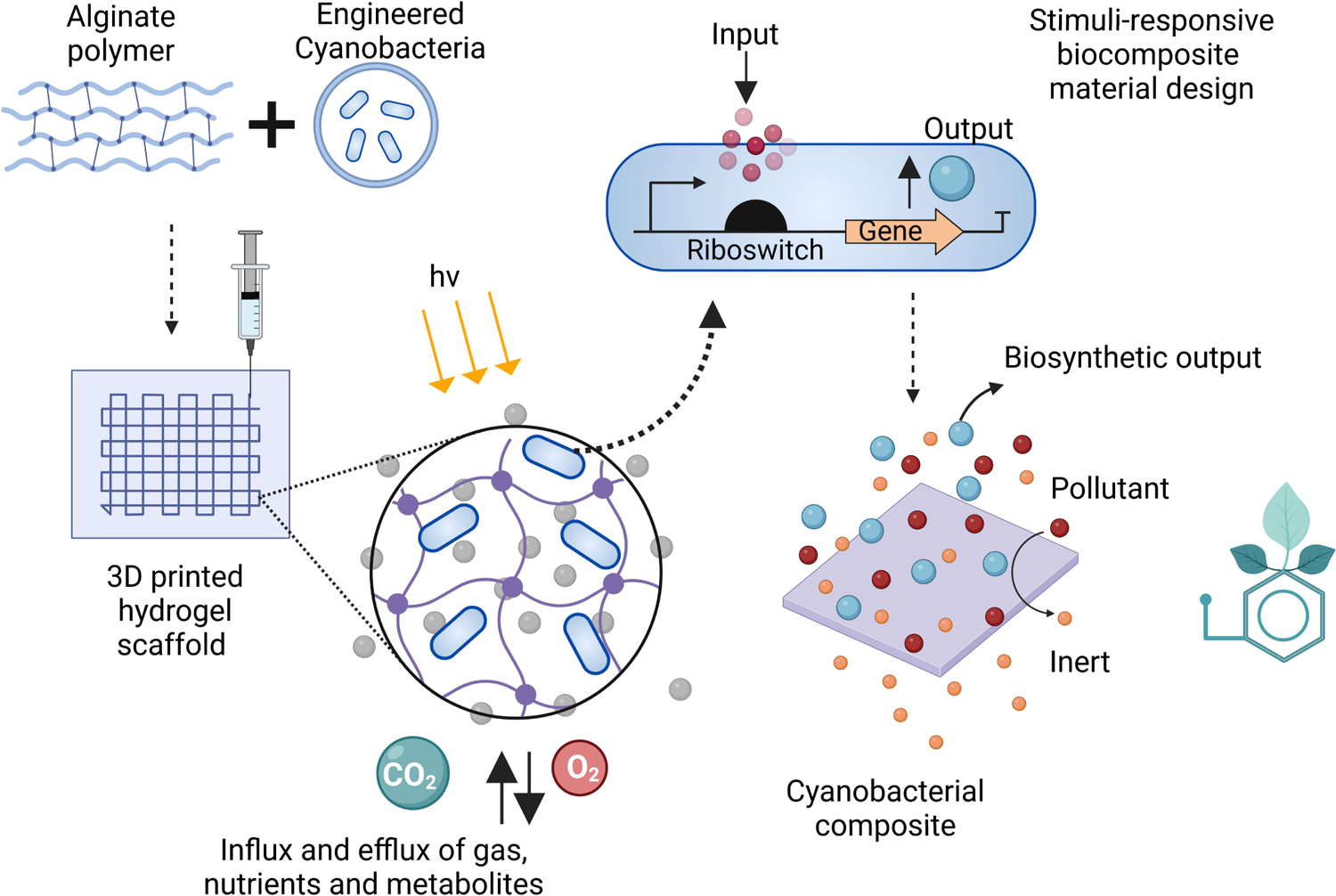
Schematic illustration of the use of engineered cyanobacteria for creating stimuli-responsive living materials used in this study. Genetically engineered strains of the obligate photoautotroph *S. elongatus* were immobilized in a matrix of the natural polymer alginate. The chosen polymer scaffold can be swollen in aqueous conditions, allows for the transport of nutrients and gases within the matrix and provides a suitable microenvironment for embedded cyanobacterial growth and the induction of gene expression by external cues.

### 3D printing of *S. elongatus* cells

To create complex geometric cellular scaffolds, we fabricated cell-laden polymeric gels using direct ink write (DIW) additive manufacturing techniques. Along with the transparency and porosity of the hydrogel material, another critical advantage of using a gel-based ink is the ability to protect the cells from mechanical shear stress during the extrusion process. This strategy minimizes damage to cell membranes and enhances cell viability in the printed patterns^38^.

Identifying a biocompatible and printable polymer is critical when encapsulating microorganisms in any 3D-printed matrix. Initial optimization experiments were conducted to identify a suitable material for encapsulating cyanobacterial cells that would sustain the growth and viability of the cells at ambient temperatures and light levels. Several natural and synthetic polymers capable of forming hydrogels were tested and our chosen candidate was an alginate hydrogel. The transparency of the hydrogel allows for light penetration, while the presence of relatively large pores within the matrix facilitates efficient diffusion of gas and nutrients^39^, and a capacity for water retention makes alginate an excellent candidate for photosynthetic ELMs. Alginate is a natural polysaccharide from seaweed composed of β-D-mannuronic acid and α-L-guluronic acid that is physically cross-linked using divalent cations^40^. The optimal condition for alginate to be crosslinked and printable at room temperature was determined to be a mixture of 4% w/v alginate solution with a 50 mM CaSO_4_ slurry suspension in a 2:1 ratio, resulting in a final 2.67% w/v alginate solution in 16.67 mM CaSO_4_. A crucial step in DIW printing is obtaining a material with suitable viscosity such that the printed structure can retain its shape post-printing^41^. This property was achieved by adding a solution of 100 mM CaCl_2_ to the hydrogel scaffold post-printing for 15 min to strengthen and stabilize the gel structure. The ink formulation was stable enough to retain its shape for various patterns, consisting of 5-17 layers of bioink (**Figure S1, Table S1**). For hydrogels embedded with cyanobacterial cells, the printed patterns were incubated under light in BG-11 medium post-printing to promote growth. To achieve an optimal gel formulation allowing for both 3D bioprinting and viability of the biological component, we utilized the same two-stage crosslinking strategy to prepare our gel-based bioink, producing a pattern with high structural integrity.

### Mechanical properties of hydrogels

Rheological measurements were carried out to test the mechanical properties of the 3D printed hydrogel with and without wild-type (WT) *S. elongatus* cells. The viscoelastic moduli with increasing oscillatory strain in both the unloaded gel (printed alginate hydrogel without cells) and the bioink (cell-laden gel) exhibited similar storage modulus values at lower oscillatory strain (**Figure S2a)**. Additionally, we observed a shift in crossover frequencies between the unloaded gel and the bioink samples (**Figure S2a**); the viscosity of the bioink was higher than the virgin alginate inks (**Figure S2b**). The gels demonstrated a shear-thinning behavior, a key requirement for DIW-based printing. While highly viscous inks are favorable for increasing print resolution, too high of a viscosity can result in clogging of the print needle and reduced cell viability. In this study, needle clogging was observed when the alginate concentrations were greater than ∼3% w/v, highlighting the necessity of a balance between printability, hydrogel integrity, and cell growth and viability. Furthermore, storage modulus (G′), loss modulus (G′′), and viscosity are influenced by factors including alginate concentration, molecular weight, crosslinker concentration (at both stages of printing), degree of crosslinking, and concentration of cells. Although we were unable to quantify the precise number of cells within the hydrogel matrix at any given timepoint due to experimental limitations, we observed that the presence of cells increases the storage modulus of the hydrogel between an oscillation strain range of ∼ 1-10 %.

### Growth and viability of *S. elongatus* cells in alginate hydrogels

Over seven days of growth, the biomass of WT *S. elongatus* cells within various designs of hydrogel constructs became visibly denser (as indicated by dark green coloration); however, designs that had low surface area to volume ratios, such as a disk design (2 cm in diameter), resulted in a decrease in biomass towards the center of the hydrogel, suggesting a limitation of gas and/or nutrient exchange (**Figure 2a**). For this reason, subsequent experiments were conducted using hydrogels printed in a 29 × 29 mm grid configuration to achieve a higher surface area to volume ratio to enhance mass transport. The grid-patterned hydrogel resulted in a greater density of biomass per unit volume hydrogel, highlighting the utility of additive manufacturing to impact geometry, and hence, the viability of the ELMs. Field emission scanning electron microscopy (FESEM) images of an unloaded gel illustrate the porous nature of the hydrogel scaffold (**Figure 2b (i & ii)**), with apparent pore sizes ranging from less than 40 μm to over 60 μm. Micrographs of hydrogels printed with bioink illustrate a combination of individual cells and colonies of cells adhered to the hydrogel surface (**Figure 2b (iii & iv)**). Given an average *S. elongatus* cell size of approximately 2 μm, and that the pore sizes observed in the biocomposite hydrogel often exceeded 60 μm, the pore diameter is likely not a limitation to the growth of cyanobacteria within the matrix. To visualize the proportion of viable to dead cells within the bioprinted matrix, the resultant hydrogels were imaged using confocal microscopy for chlorophyll autofluorescence and cell death using SYTOX Blue, a blue nucleic acid stain that penetrates cells with damaged cell membranes (**Figure 2c**). Red fluorescent patches, indicative of healthy colonies of cyanobacteria, were visible throughout the hydrogel over seven days of growth, whereas blue patches indicative of dead cells were distributed heterogeneously in the hydrogel.

**Figure 2.**
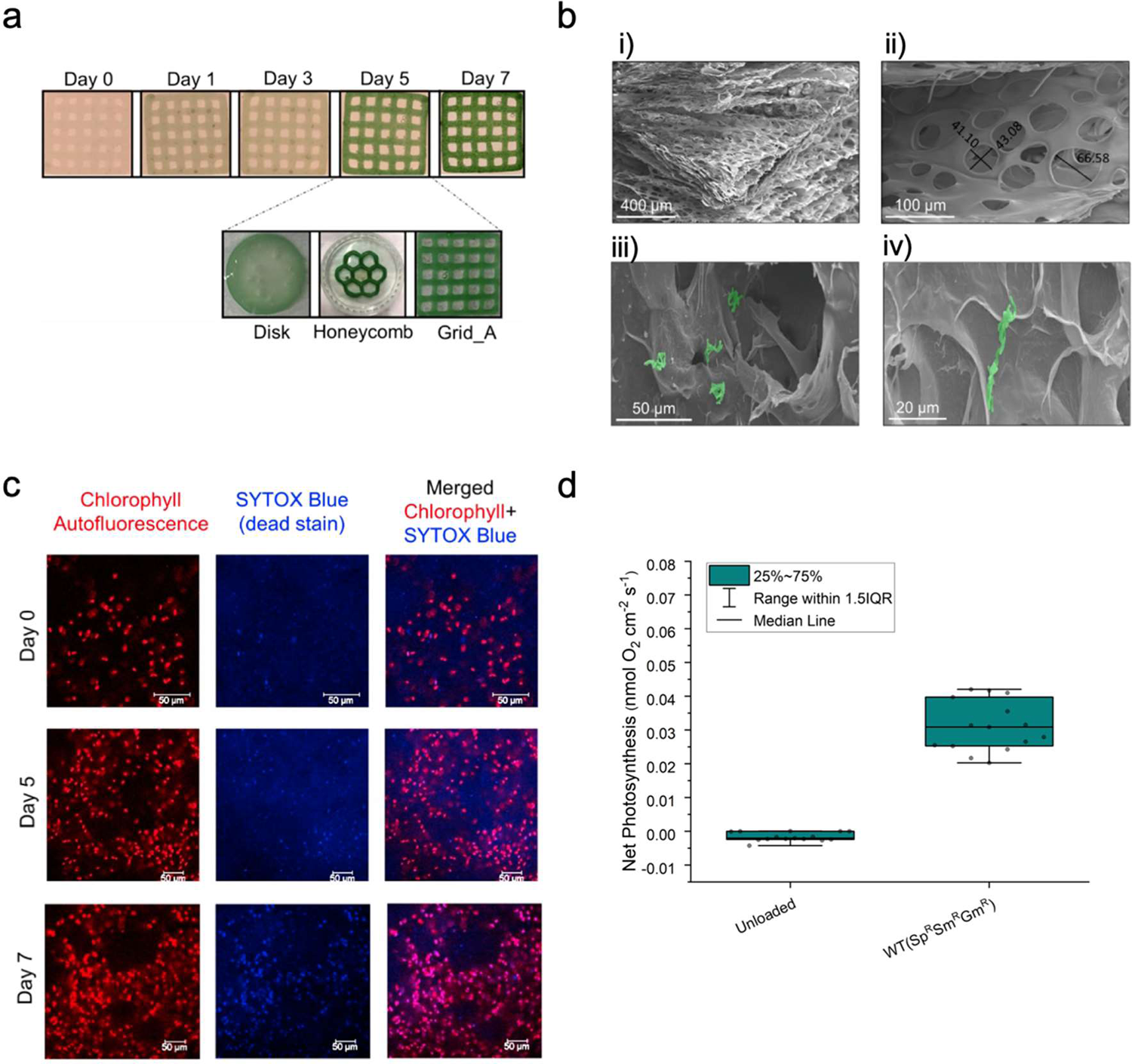
a) Top: Growth of WT *S. elongatus* within a 3D printed grid pattern over 7 days. Bottom: Images of 5-day-old hydrogels containing WT cells printed on a disk, honeycomb, and grid_A geometry. The dimension details of these different patterns are described in Supplementary Table S1. b) FESEM images of unloaded hydrogels (i & ii), and hydrogels containing WT *S. elongatus* cells (iii & iv). *S. elongatus* cells are highlighted in false color green. c) Confocal microscopy images of chlorophyll autofluorescence and SYTOX blue staining of a hydrogel containing WT *S. elongatus* cells after printing and 0, 5, and 7 days of growth. d) Box plot of net photosynthesis at an incident downwelling irradiance of 80 μmol photon m^-2^ s^-1^ for an unloaded hydrogel and an antibiotic resistant *S. elongatus* strain [WT(Sp^R^Sm^R^Gm^R^)] encapsulated in the hydrogel.

### Photosynthetic activity of living hydrogels

To assess the photosynthetic activity of the living hydrogels, the O_2_ microenvironment and O_2_ turnover were measured in light and darkness using O_2_ microsensors. O_2_ evolution was detected from an antibiotic-resistant *S. elongatus* strain WT(Sp^R^Sm^R^Gm^R^), which is used as a control for engineered strains described in later sections (**Table S4**), embedded in hydrogels (**Figure S3a**). O_2_ microsensors revealed an actively photosynthesizing and hyperoxic microenvironment on the surface of the living hydrogels (**Figure S3b**). For the hydrogel containing WT(Sp^R^Sm^R^Gm^R^) cells, mean O_2_ concentration adjacent to the hydrogel surface was 346 ± 4.9 μM SE; (Tukey post hoc p = <0.01) at an incident irradiance of 80 μmol photons m^-2^ s^-1^. Net photosynthesis (**Figure 2d)** observed for the WT(Sp^R^Sm^R^Gm^R^) strain was 0.031 nmol O_2_ cm^-2^ s^-1^ (Tukey post hoc, p < 0.01). No oxygen evolution was detected from the unloaded hydrogel; however, slight respiratory activity was detected indicative of the presence of low levels of opportunistic bacteria. During darkness hydrogels encapsulating the WT(Sp^R^Sm^R^Gm^R^) strain showed limited respiratory activity and O_2_ concentrations were close to ambient seawater values (**Figure S3e**). Measurements of variable chlorophyll a fluorimetry were performed to determine the photosynthetic efficiency of the living hydrogels over a range of irradiance regimes. For the WT(Sp^R^Sm^R^Gm^R^) *S. elongatus* strain, photosynthesis was saturated between 200 – 250 μmol photons m^-2^ s^-1^ with an irradiance at onset of saturation (E_k_= P_max_/α) of about 85 μmol photons m^-2^ s^-1^, indicative of low light adaptation (**Figure S3d**). Electron transport above 250 μmol photons m^-2^ s^-1^ was fully inhibited (**Figure S3f**).

While hydrogels containing *S. elongatus* could grow from low initial cell densities under low to moderate light, growth was arrested in samples grown under irradiances greater than 300 μmol photon m^-2^ s^-1^. Photosynthesis vs irradiance curves support saturation of photosynthesis at similar irradiance levels for these low-light adapted hydrogels (**Figure S3d**). Because solar radiation levels on the earth’s surface can reach approximately 2000 μmol photons m^-2^ s^-1^, freshly printed biocomposites will likely suffer in natural high-light environments. However, aquatic environments such as lakes and rivers often contain large concentrations of colored dissolved organic matter that rapidly attenuate light with depth^42^, providing a hospitable environment for the ELMs presented here.

### Regulation of YFP expression in hydrogels

To investigate whether cells grown within the hydrogel could respond to an external chemical stimulus, *S. elongatus* was transformed with plasmids pAM4909, pAM5027, and pAM5057 to yield strains YFP^+^ (constitutive-expression YFP positive control), YFP^-^ (negative control), and RiboF-YFP^+^ (riboswitch-regulated YFP expression), respectively (**Table S2, S4**). A schematic of the RiboF-YFP^+^ genetic circuit is illustrated in Figure 3a; the engineered *S. elongatus* strain responds to the chemical inducer theophylline from the surrounding environment by a conformational change in mRNA bearing Riboswitch-F, which regulates translation of the YFP reporter protein. The YFP^+^ construct contains a constitutive *con*II promoter driving the expression of YFP. After treatment with 1 mM theophylline in dimethyl sulfoxide (DMSO) or a 1% DMSO vehicle control for 24 hours, YFP expression was determined qualitatively within the hydrogel using fluorescence microscopy (**Figure 3b**). No YFP fluorescence was observed from hydrogels containing the YFP^-^ strain. Constitutive YFP fluorescence was observed from hydrogels containing the YFP^+^ strain in medium containing either 1 mM theophylline or 1% DMSO. For hydrogels containing the RiboF-YFP^+^ strain, growth in 1 mM theophylline produced similar fluorescence levels to hydrogels containing the YFP^+^ control strain, while no YFP fluorescence was detected in hydrogels supplemented with 1% DMSO. These data demonstrate that the cells embedded within the hydrogel can respond to an external chemical stimulus and be engineered for tight regulation of expression.

**Figure 3.**
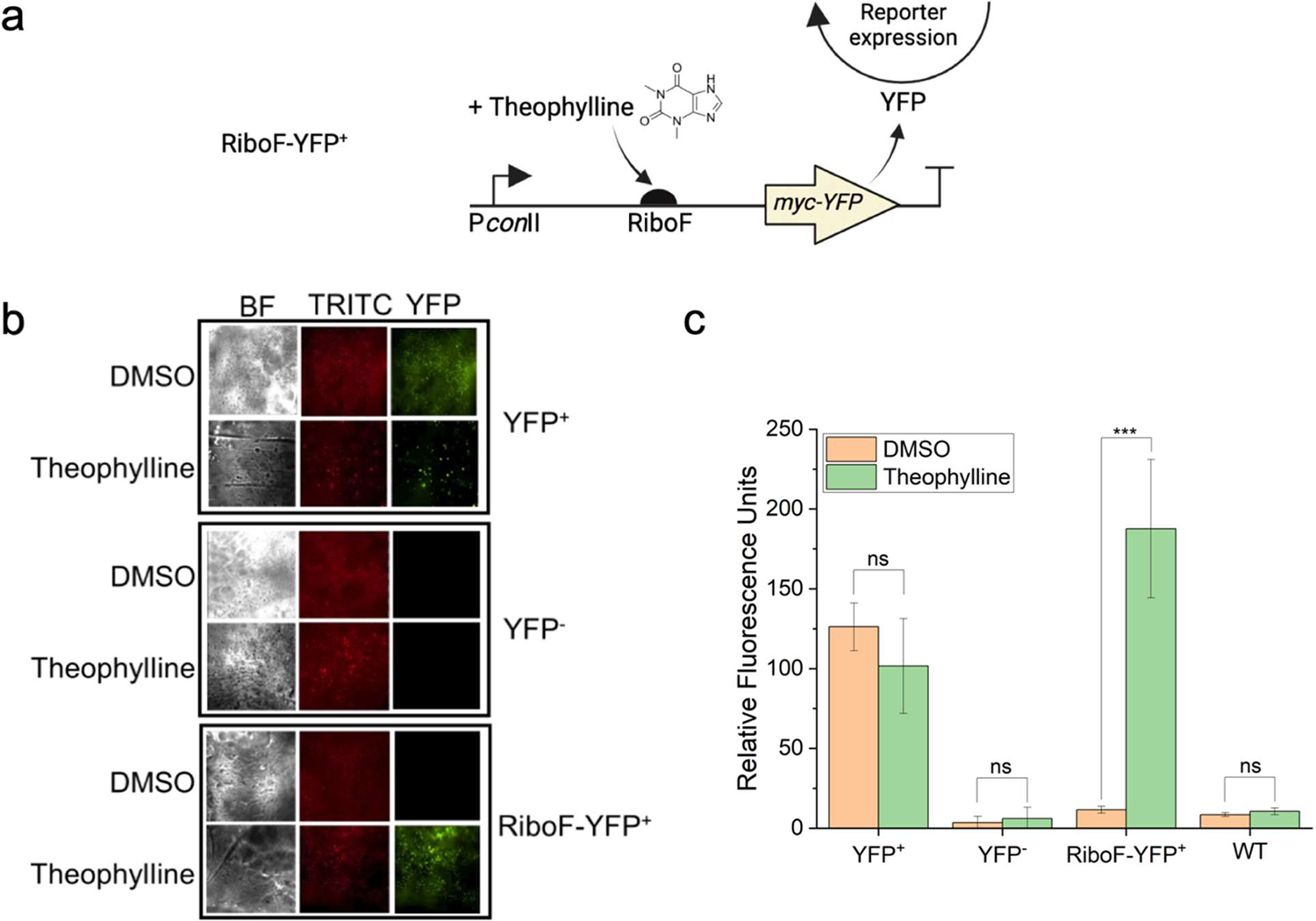
a) Schematic overview of the genetic circuit used in the construction of RiboF-YFP^+^ strain. b) Fluorescence microscopy images of hydrogels containing cells expressing a YFP reporter. Hydrogels containing the YFP^+^, YFP^-^, and RiboF-YFP^+^ strains were supplemented with either 1% DMSO vehicle control or 1 mM theophylline. Images are shown for the hydrogels under brightfield (BF), TRITC, and YFP channels. c) YFP fluorescence measured from the BG-11 medium each ELM structure was incubated in.

To assess whether viable engineered cells may have leaked from the hydrogel into the surrounding medium, media from 5-day old hydrogels containing the YFP^-^, YFP^+^, or RiboF-YFP^+^ strains were incubated with 1 mM theophylline or 1% DMSO for 24 hours and YFP fluorescence was quantified with a fluorescence plate reader. An approximately 16-fold increase in fluorescence was detected in supernatants from the RiboF-YFP^+^ samples induced with theophylline compared to the uninduced sample (**Figure 3c**). No significant difference was observed between induced and uninduced samples of supernatants from all other hydrogels. The presence of viable cells in the surrounding media may be attributed to a low level of diffusion of cells via mechanical agitation in an aqueous solution throughout the hydrogel and at the hydrogel-medium interface, resulting in a slow release of cells.

### Construction of riboswitch-induced laccase strains and ABTS-Laccase activity testing for 3D printed constructs

To test the utility of the riboswitch regulatory mechanism for expression of a functional enzyme, plasmid construct pAM5826 (**Table S2, S4**) was created containing a theophylline-responsive riboswitch upstream of the *cotA* gene in pAM5825 and used to insert the cassette into the *S. elongatus* chromosome to produce strain RiboF-Laccase^+^. **Figure 4a** illustrates a schematic of the RiboF-Laccase^+^ genetic circuit.

**Figure 4.**
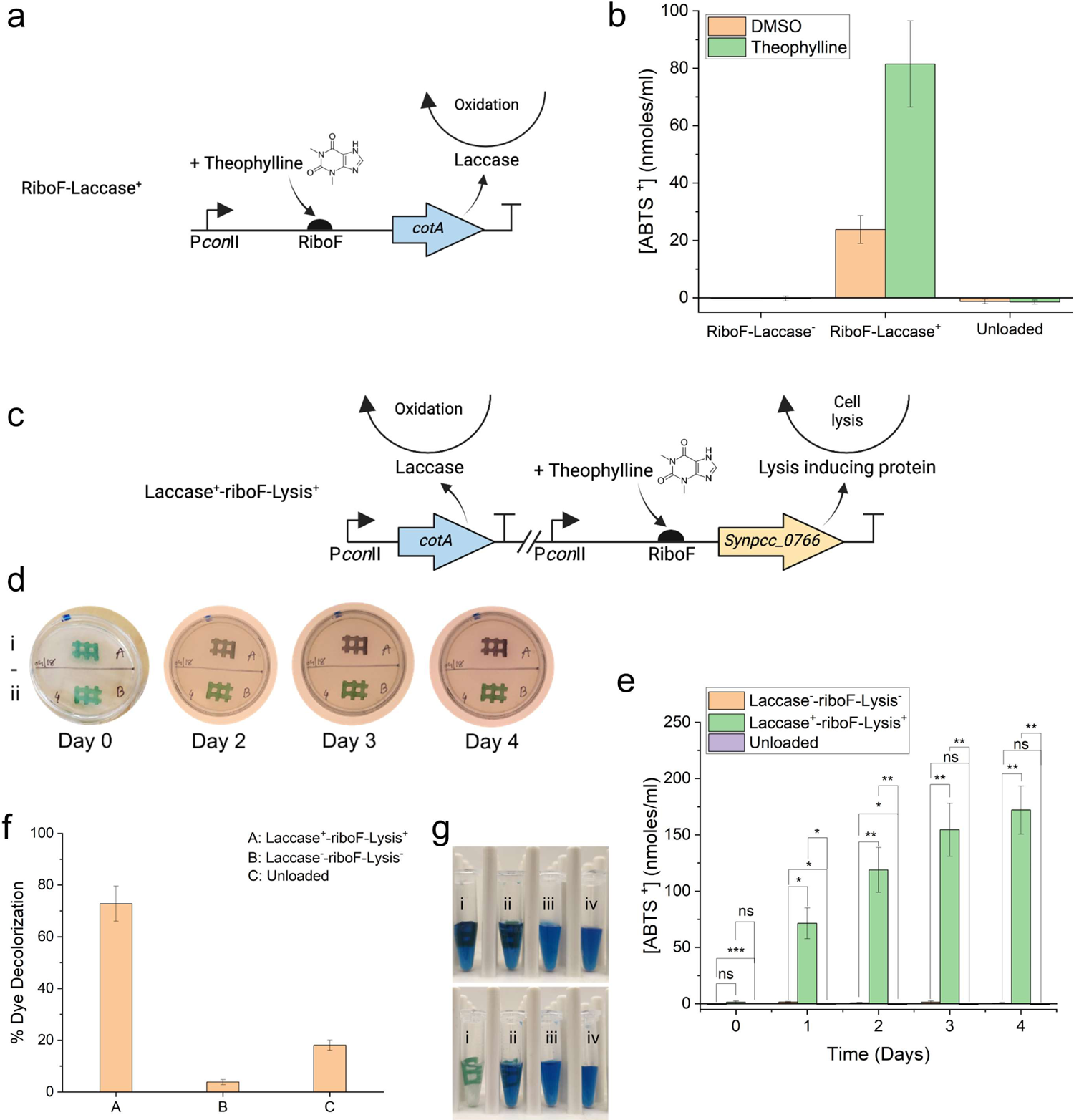
a) Schematic overview of the genetic circuit used in the construction of the RiboF-Laccase^+^ strain. b) Oxidation of ABTS in reaction buffer by hydrogels containing strains RiboF-Laccase^-^ control, RiboF-Laccase^+^, or unloaded hydrogel pre-incubated with either 1% DMSO or 1 mM theophylline. c) Schematic overview of the genetic circuit used in the construction of Laccase^+^-riboF-Lysis^+^ strain. d) Visual assay of hydrogel ABTS oxidation activity over four days incubation in standard growth conditions. i) Subsection of a hydrogel containing strain Laccase^+^-riboF-Lysis^+^. ii) Subsection of a hydrogel containing the Laccase^-^-riboF-Lysis^-^ control strain. e) Time-course of the oxidation of ABTS in reaction buffer after the addition of hydrogels containing either the Laccase^-^-riboF-Lysis^-^ control strain, Laccase^+^-riboF-Lysis^+^ strain, or an unloaded hydrogel control. f) Percent decolorization of indigo carmine (0.1 mg/mL in BG-11) by hydrogels printed with either Laccase^-^-riboF-Lysis^-^ or Laccase^+^-riboF-Lysis^+^ strains or unloaded hydrogel after incubation for 10 days. g) Images of hydrogels at incubation time 0 (top) and after 10 days of incubation (bottom). (Roman numerals indicate the addition of i) hydrogel printed with Laccase^+^-riboF-Lysis^+^, ii) hydrogel printed with Laccase^-^-riboF-Lysis^-^, iii) unloaded hydrogel, iv) BG-11 with dye only.)

The surrounding solution of the hydrogel ELMs was tested for oxidative activity of the laccase enzyme against the substrate 2,2′-Azino-bis-(3-ethylbenzothiazoline-6-sulfonic acid) (ABTS). Briefly, after 5 days of growth, hydrogels containing strain RiboF-Laccase^+^, RiboF-Laccase^-^, and an unloaded-control were transferred to fresh BG-11 medium supplemented with either 1 mM theophylline in DMSO or a 1% DMSO vehicle control and incubated for 72 hours. The hydrogels were then transferred to a buffer solution containing ABTS at a final concentration of 2 mM and incubated for 24 hours. The laccase activity was measured by quantifying concentrations of oxidized ABTS from the surrounding solution (**Figure 4b)**. No oxidized ABTS was detected in either the RiboF-Laccase^-^-containing or unloaded hydrogel samples. Levels of oxidized ABTS were 3.4-fold higher in RiboF-Laccase^+^ samples with theophylline induction relative to uninduced samples.

While YFP fluorescence was nearly undetectable in uninduced RiboF-YFP^+^ strains, moderate levels of laccase activity were detected in uninduced RiboF-Laccase^+^ samples. This difference is likely because of assay sensitivity due to the high activity of CotA and the low enzyme level required to oxidize ABTS. For future applications requiring tighter regulation of protein products, incorporating multiple riboswitches or pairing transcriptional regulatory systems with different riboswitches could be used to fine-tune expression.

### Construction of constitutive laccase-expressing and inducible cell death strains

To create living hydrogels suitable for bioremediation, a strain capable of constitutively producing the laccase enzyme was constructed with an additional feature of inducible cell death. In response to an observed basal level of cell leakage from the hydrogels over time, the system was engineered for inducible cell death as a mechanism to minimize the potential for contamination and biofouling by the ELM. A neutral-site 2 (NS2) chromosome-integration plasmid (pAM5825) was constructed that encodes a laccase enzyme (CotA) expressed from the constitutive *con*II promoter (**Table S2, S4**). The construct was inserted into the *S. elongatus* chromosome in addition to a construct from plasmid pAM5829 (**Table S2**), containing a *con*II promoter–theophylline-responsive riboswitch construct upstream of the native *S. elongatus* gene Synpcc7942_0766, creating strain Laccase^+^-riboF-Lysis^+^ and printed in hydrogels in tandem with a corresponding control strain Laccase^-^-riboF-Lysis^-^. **Figure 4c** illustrates a schematic of the Laccase^+^-riboF-Lysis^+^ genetic circuit.

Concentrations of O_2_ at the hydrogel surface and oxygen evolution and respiration rates of the hydrogels embedded with the Laccase^+^-riboF-Lysis^+^ were similar to hydrogels embedded with the WT(Sp^R^Sm^R^Gm^R^) strain (Figure S3a-e; see supplementary information text for details), and genotypic characterization and expression of the 52-kDa CotA enzyme were confirmed (**Figure S4a, b**).

### ABTS-Laccase activity test for 3D printed Laccase^+^-riboF-Lysis^+^ and control Laccase^-^-riboF-Lysis^-^ patterns

The enzyme activity was tested against ABTS both within the ELM hydrogel and from the surrounding solution. The hydrogels were first separated from their surrounding BG-11 media and submerged in a reaction buffer mixture with 2 mM ABTS. Following 1 hour of incubation, a set of hydrogels containing Laccase^-^-riboF-Lysis^-^ and Laccase^+^-riboF-Lysis^+^ saturated with ABTS solution were removed, placed on sterile agar LB plates, and left to incubate for 4 days (**Figure 4d**). Over the 4-day period, a purple coloration developed in the Laccase^+^-riboF-Lysis^+^ hydrogel indicative of the oxidation of ABTS, while the hydrogel containing the Laccase^-^-riboF-Lysis^-^ strain remained the natural green hue produced by cyanobacterial pigments that was observed in all patterns at time 0.

To quantify the laccase activity of the ELMs in the surrounding solution, a set of hydrogels containing Laccase^-^-riboF-Lysis^-^, Laccase^+^-riboF-Lysis^+^, and unloaded gels were submerged in a buffer solution containing ABTS and incubated for 4 days with daily sampling for quantification of the oxidation of ABTS in the reaction buffer (**Figure 4e**). After 24 hours, 71.5 ± 13.6 nmol/ml of ABTS was oxidized by the Laccase^+^-riboF-Lysis^+^ hydrogel. A small but significant level of activity against ABTS was observed in the Laccase^-^-riboF-Lysis^-^ relative to the control unloaded gel. Over the course of 4 days of incubation, the level of oxidized ABTS from the hydrogel embedded with Laccase^+^-riboF-Lysis^+^ increased to 172.0 ± 21.3 nmol/mL, which was significantly greater than either the Laccase^-^-riboF-Lysis^-^**-**containing or unloaded gels (*P* value < 0.01). The oxidation of the ABTS substrate to its cationic radical form was also evident from the dark green coloration observed in the hydrogels bearing strain Laccase^+^-riboF-Lysis^+^ over time (**Figure S5**).

Additionally, the laccase activity of the supernatant of media surrounding hydrogels was measured from samples taken five days post-printing, to assess whether the CotA enzyme escapes from the hydrogel, either by means of secretion or cell lysis within the gel (**Figure S6**). The activity observed from the supernatant surrounding the Laccase^+^-riboF-Lysis^+^ containing hydrogels was significantly greater than either the Laccase^-^-riboF-Lysis^-^ containing or unloaded gels. However, a low level of ABTS oxidation was observed in supernatants from hydrogels embedded with Laccase^-^-riboF-Lysis^-^ relative to unloaded hydrogels at each timepoint, indicating that the secretion or lysis products of the control Laccase^-^-riboF-Lysis^-^ strain also confer some level of oxidative activity against the substrate ABTS, albeit a reduced level relative to the hydrogels embedded with Laccase^+^-riboF-Lysis^+^.

### Indigo carmine oxidation activity for 3D printed Laccase^+^-riboF-Lysis^+^ and control Laccase^-^ -riboF-Lysis^-^ patterns

To assess whether hydrogels containing the engineered strains of Laccase^+^-riboF-Lysis^+^ can serve as a tool for bioremediation, the hydrogels were assayed for their ability to decolorize a common textile dye pollutant, indigo carmine, a capability previously demonstrated for a heterologously-expressed laccase^35^. Five-day old hydrogels containing either Laccase^+^-riboF-Lysis^+^, Laccase^-^-riboF-Lysis^-^, or an unloaded-control were submerged in a BG-11 solution containing 0.1 mg/ml of indigo carmine. The decolorization of indigo carmine was subsequently monitored over the course of 10 days and quantified by proxy of absorption at 612 nm, the absorption maximum of indigo carmine in the visible spectrum (**Figure 4f, 4g**). The hydrogel samples embedded with Laccase^+^-riboF-Lysis^+^ had an average decolorization of 72.8 ± 6.8 %, while the Laccase^-^-riboF-Lysis^-^ and unloaded hydrogel had decolorization averages of 3.8 ± 1.0 % and 18.1 ± 2.0 %, respectively. The percent dye decolorization in hydrogels containing Laccase^-^ -riboF-Lysis^-^ and unloaded hydrogels is due to the adsorption of indigo carmine by the alginate polymer matrix, as has been previously reported for other substrates^43,44^, and is observable as the blue hue of unloaded hydrogels removed from the solution (**Figure S7**).

The hydrogel embedded with Laccase^+^-riboF-Lysis^+^ successfully decolorized ∼70% of the indigo carmine over the course of 10 days with the added benefit of being physically removable from the system without requiring energy-intensive steps to clarify a liquid culture, such as centrifugation. The unloaded hydrogel also produced a significant level of apparent indigo carmine decolorization due to the absorption of indigo carmine by the hydrogel matrix itself. Interestingly, the unloaded hydrogel consistently had a greater level of decolorization due to non-specific adsorption of the dye molecule than did hydrogels containing the Laccase^-^-riboF-Lysis^-^ control strain.

### Inducible cell death of cyanobacteria

When designing ELMs, consideration of the unintended consequences of contamination from the biological component into the surrounding environment should be taken into consideration. The Synpcc7942_0766 gene, when overexpressed, results in the excision of a defective prophage from the *S. elongatus* genome^45^, resulting in cellular lysis (**Figure 5a**). Hydrogels containing Laccase^+^-riboF-Lysis^+^ or Laccase^-^-riboF-Lysis^-^ (control) strains were transferred to fresh medium after 5 days of initial growth and incubated for an additional 12 days in fresh growth media supplemented with either 1 mM theophylline in DMSO, 1% DMSO, or medium only (**Figure 5b**). After 12 days of growth, visual inspection indicated that accumulation of biomass in the surrounding medium was absent in the flask containing the Laccase^+^-riboF-Lysis^+^ strain supplemented with 1 mM theophylline. The timescale of response to induction of cell death in the surrounding medium was assessed by supplementing fresh hydrogel supernatants containing cells of either Laccase^+^-riboF-Lysis^+^ or Laccase^-^-riboF-Lysis^-^ strains with either 1 mM theophylline or 1% DMSO and monitoring growth of the cultures for four days (**Figure 5c**). Within 48 hours, ectopic induction of Synpcc7942_0766 in the Laccase^+^-riboF-Lysis^+^ strain supplemented with 1 mM theophylline resulted in a decrease in OD_750_, initially visible as clumps of senescent cell matter that continued to break down to produce a pale hazy medium, indicating death of the culture (**Figure 5c, 5d**). Cultures of Laccase^+^-riboF-Lysis^+^ supplemented with 1% DMSO and Laccase^-^-riboF-Lysis^-^ supplemented with either 1 mM theophylline or 1% DMSO continued to grow at similar rates to one another, demonstrating the utility of engineered inducible cell death in an ELM as a mechanism of targeted reduction of biological contamination by the material.

**Figure 5.**
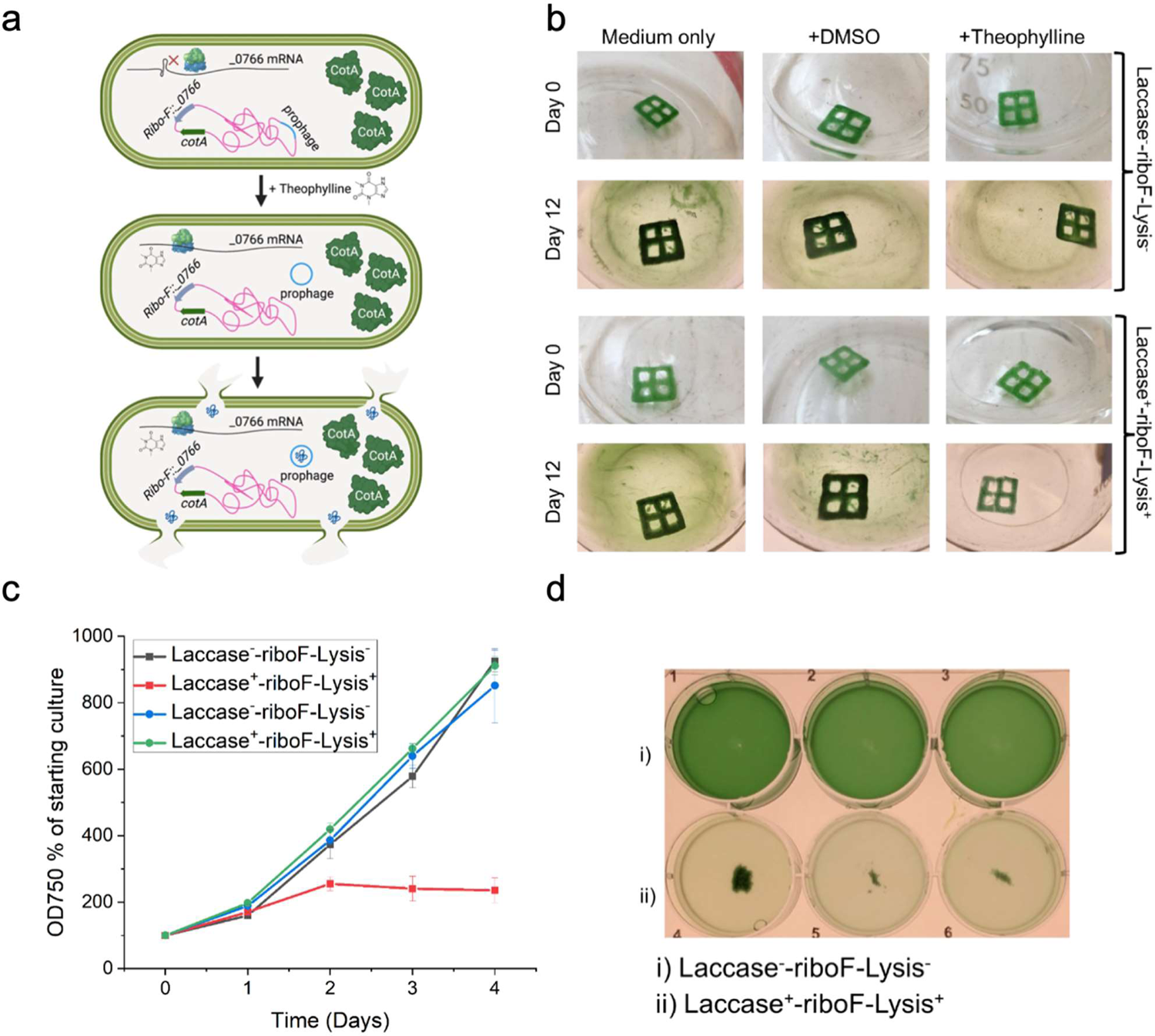
Inducible cell death of strain Laccase^+^-riboF-Lysis^+^. a) Schematic of constitutive expression of CotA, and riboswitch conformation change on with the addition of theophylline, leading to the overexpression of Synpcc7942_0766, prophage excision, and subsequent cell death. b) Day 0 and day 12 images of the hydrogels containing Laccase^-^-riboF-Lysis^-^ control and Laccase^+^-riboF-Lysis^+^ strains supplemented with either medium only, 1% DMSO, or 1 mM theophylline. c) Growth curves of the Laccase^-^-riboF-Lysis^-^ control and Laccase^+^-riboF-Lysis^+^ strains with the addition of either 1 mM theophylline (square) or 1% DMSO (circle) in solution. d) Image of liquid cultures of Laccase^-^-riboF-Lysis^-^ and Laccase^+^-riboF-Lysis^+^ strains three days post supplementation with 1 mM theophylline.

Riboswitches are increasingly being used for applied bioengineering due to the simplicity of ligand-aptamer interactions^13,46,47^. The use of riboswitches to regulate protein expression within hydrogels was successful for YFP, laccase, and the lysis-inducing protein and should function well in future biocomposites that contain alternative cyanobacterial or algal strains. We attribute the success of regulation by riboswitches to the diffusion of the small molecule theophylline throughout the porous alginate hydrogel.

### Outlook

We optimized the composition of an alginate-based hydrogel for 3D printing and viability of the cyanobacterium *S. elongatus*, and engineered several strains of the cyanobacterium to be stimulus-responsive and produce functional outputs when embedded within the hydrogel. Previously, hydrogels constructed with cyanobacteria as a biological component^36,48,49^ have been limited to wild-type strains. In this study, we leveraged genetic toolkits for synthetic biology to produce photosynthetic ELMs with functional outputs and tailored regulatory circuits. The use of riboswitches to regulate gene expression within the living material yielded stimulus-responsive materials with the capacity for dye-decolorization and inducible cell death to prevent cellular contamination in the environment.

While the photosynthetic ELMs designed in this study have functional outputs suitable for bioremediation, numerous improvements could make the material more appropriate for use outside of the laboratory setting. Notably, improvements in enzymatic activity per unit volume of hydrogel will be necessary to move from the laboratory scale to large-scale applications, such as bioremediation in lakes or water treatment plants. One strategy for the improvement of activity from the material could be the modification of protein products to be tagged for secretion from the cell resulting in higher concentrations in solution and allowing for the production of bioproducts that can be toxic to the cells at high intercellular concentrations. Knowledge of mechanisms and systems for the secretion of larger recombinant proteins in cyanobacteria such as *S. elongatus* PCC 7942 will greatly benefit the output potential of cyanobacterium-based ELMs. Secondly, careful consideration should be paid to the influence of materials containing biological components on the natural environment, such as biofouling and subsequent effects on ecological systems. The ELMs presented here address this issue by incorporating a mechanism for inducible cell death by the addition of the small molecule theophylline. While theophylline is a natural product present in small quantities in tea and cocoa, it is not native to aquatic systems and its use in the natural environment could pose risk of unintended chemo-ecological consequences. Alternative regulatory circuits with more compatible inductive signals could be engineered for use in the natural environment. For example, spectrally responsive photoreceptors or temperature-sensitive signaling transduction pathways could allow for diurnal or seasonal cell death of biological contamination.

The ELM presented here serves as a proof-of-concept for the engineering and incorporation of cyanobacteria into a polymeric matrix to produce stimuli-responsive materials with functional outputs of real-world utility. Utilizing cyanobacteria as the biological component, these ELMs fundamentally depend on only light, CO_2_ as a carbon source, and minimal nutrients for survival. While the biocomposites demonstrated here have been designed for use in bioremediation, the functional outputs of cyanobacterial ELMs could be engineered to be broadly impactful.

## Materials and Methods

### Cultivation of cyanobacteria

Wild-type control and genetically engineered *Synechococcus elongatus* PCC 7942 cyanobacterial cells were cultivated in BG-11 medium^50^ in conical flasks and grown at 30° C while shaking at 120 rpm under continuous illumination of 75 μmol photons m^-2^ s^-1^. BG-11 was supplemented with 2 μg/ml spectinomycin, 2 μg/ml streptomycin, and/or 2 μg/ml gentamycin for selection as required. Unless otherwise noted, cells were grown for 5-6 days from an initial OD_750_ of 0.05 to mid-log phase (OD_750_ = 0.5-0.6) prior to the preparation of bioinks.

### Preparation of alginate ink and bioink

Alginate hydrogels were prepared using a mixture of sodium alginate with CaSO_4_ and CaCl_2_ as crosslinking agents at optimized concentrations. In brief, a 4% w/v sodium alginate solution was prepared by dissolving sodium alginate powder (Sigma-Aldrich) in autoclaved deionized water. The solution was continuously stirred overnight at room temperature, achieving a clear, viscous solution. Stock solutions of CaSO_4_ and CaCl_2_ were prepared in autoclaved deionized water at concentrations of 50 mM and 100 mM, respectively. All solutions were stored at 4 °C until use. A hydrogel ink solution was prepared for printing by mixing the alginate solution and CaSO_4_ in a 2:1 ratio. The slurry suspension of 50 mM CaSO_4_ was mixed thoroughly (25-30 times) with alginate solution using an in-house fabricated 3-way syringe mixer and then transferred to a 3 cc syringe for 3D printing. The ink in the 3 cc syringe was placed within a 50 mL falcon tube and centrifuged at 4000 rpm for 10 min to remove any air bubbles prior to printing.

Bioinks were prepared with cells from cultures at an OD_750_ of 0.5-0.6. The cells were collected by centrifugation at 3500 x g for 5 min. The pellet was mixed in the plotting paste using the 3-way syringe mixer (**Figure S8**). All glassware used in the process was pre-sterilized by autoclaving and all accessories required for cell mixing and 3D printing were pre-sterilized by UV sterilization. Both the ink and the bioink were prepared fresh prior to each experiment.

### 3D printing of alginate ink and bioink

CaSO_4_ was used to partially crosslink the alginate chains to make the gel suitable for Direct-Ink-Writing (DIW) printing. Various combinations of alginate solutions (0.5-5% w/v) and CaSO_4_ (25-100 mM) concentrations were tested at different pressures (50-350 kPa) using steel blunt end syringe needles (22, 27-, and 32-gauge sizes) to optimize conditions for printability. The homogeneous mixture that gave the most stable structure at a given needle size and pressure was used for printing the bioink. In brief, the ink with or without cyanobacterial cells was loaded in a 3 cc printing syringe barrel, equipped with a 27-gauge needle. The ink was 3-D printed with a pre-designed geometry using the Inkredible+ Bioprinter (Cellink, USA). The printing parameters were as follows: pressure ∼200 kPa, infill density 50%, fill pattern rectilinear, and speed 80 mm/sec. All prints were produced in ambient conditions at room temperature. The printed geometries were designed using CAD software and converted to Gcode using the Cellink Heartware software. Ink without *S. elongatus* cells was used as a cell-free control hydrogel in all experiments.

Post printing, the multilayered 3D-printed structures were immersed in 100 mM CaCl_2_ for 15 min to allow the patterns to be fully crosslinked and then incubated in 47 mm Petri dishes containing 5 ml BG-11 medium supplemented with antibiotics. Unless otherwise mentioned, all the scaffolds with either WT or engineered strains were kept in an incubator at an irradiance level of 20 μmol photons m^-2^ s^-1^ and 28 °C for initial growth prior to experimentation.

### Rheological Characterization and Viscosity measurements

Viscoelastic moduli and viscosity measurements of the alginate ink with and without the inclusion of WT cells were carried out using a Discovery HR-30 Hybrid Rheometer (TA Instruments). Both measurements were conducted using a 20 mm parallel plate geometry in a Peltier plate setup. For amplitude sweep measurements, angular frequency was 10 rad/sec and strain ranged from 0.01 % to 1000.0 %. For viscosity, flow sweep was measured at a continuous shear rate of 0.01 s^-1^ to 1000.0 s^-1^. All characterizations were conducted at a temperature of 25 °C.

### Growth optimization of the 3D-printed WT *S. elongatus*

The growth of WT *S. elongatus* cells was monitored and optimized in different geometries of 3D-printed patterns over time. Specifically, cells encapsulated in three different printed structures (disk, honeycomb, and grid_A) were incubated in BG-11 medium with constant illumination for 7 days and were visually monitored for color density over time. Images were taken at regular time intervals.

### Cell viability of 3D-printed WT *S. elongatus*

Cell viability of WT *S. elongatus* within the 3D-printed patterns was assessed using confocal microscopy. Live/dead cell assays were performed by imaging chlorophyll autofluorescence and SYTOX Blue (SB; Fisher Scientific) stain fluorescence. 3D-printed grid_A patterns were cut to yield two interconnected hollow squares under sterile conditions at regular time intervals and incubated in 5 μL of a 1 mM SB stock solution in 1 mL of BG-11 medium for 5 min in the dark. The effective concentration of the stain was 5 μM. After incubation, the samples were washed three times with BG-11 and analyzed on a Leica Sp8 confocal microscope with lightning deconvolution and white light laser. The channel for SB detection was set at excitation 405 nm, and the emission was collected at 450–510 nm. The channel for chlorophyll autofluorescence was set at excitation 488 nm and the emission was collected at 685-720 nm. Samples from different time points were imaged to monitor cell viability.

### Field Emission Scanning Electron Microscopy

Electron microscopy images of multiple regions of freeze-dried hydrogel samples, printed with and without WT cells, were taken on an FEI Apreo FESEM (Thermo Scientific). Sample subsections were cut and stored at −80 °C for 24 hours. The samples were subsequently lyophilized to remove residual water, and the dried samples were prepared for imaging. The samples were adhered to SEM sample stubs using double-sided carbon tape and were coated uniformly with gold particles for 60 msec. Images were taken using an ETD detector at 10-20kV.

### O_2_ microsensor measurements

Clark-type O_2_ microsensors (tip size 25–50 μm, Unisense, Aarhus, Denmark) were used to measure the O_2_ microenvironment and O_2_ turnover of the living hydrogels in light and in darkness. Microsensor measurements were performed as described previously^51,52^. O_2_ sensors were mounted on a motorized micromanipulator (MU1, PyroScience GmbH, Germany) that was attached to an optical table. Cell-loaded and unloaded hydrogels were placed in a custom-made acrylic system that provided slow laminar flow at a rate of approximately 1 cm/s. O_2_ microsensor measurements were performed at the cross-section between the printed horizontal and vertical areas. Preliminary O_2_ mapping suggested that these are the areas of highest photosynthetic activity. Microsensor profiles were generated by carefully positioning the sensor at the surface of the hydrogel with the aid of a digital microscope (Dino-Lite, US). O_2_ profiles were measured from the hydrogel surface through the diffusive boundary layer into the overlying water column in steps of 50-100 μm using the motorized micromanipulator; the data are reported as means ± SE. Net photosynthesis was determined at an incident downwelling irradiance of 80 μmol photons m^-2^ s^-1^ as provided by a fiber optic halogen light source (ACE, Schott GmbH). Dark respiration was calculated as the diffusive O_2_ flux in darkness. The diffusive O_2_ flux was calculated using Fick’s first law of diffusion as described previously^51^ using a diffusion coefficient of DO_2_ = 2.2186 × 10^−5^. For each experimental treatment (unloaded hydrogel control gel, WT(Sp^R^Sm^R^Gm^R^) control gel, and Laccase^+^-riboF-Lysis^+^ gel) a total of 15 replicate measurements were performed from three printed hydrogels. The index of light adaptation (i.e., the irradiance at onset of saturation) E_k_ was calculated as E_k_ = P_max_/α; where P_max_= maximal photosynthesis rate and α = the initial slope of the light curve indicating light use efficiency.

### YFP monitoring by fluorescence microscopy from cell-laden hydrogel

YFP fluorescence from the cells within the hydrogel matrix was qualitatively measured by capturing images of hydrogel patterns on a Nikon Eclipse Ni fluorescence microscope equipped with a Nikon DS Qi2 camera at 20x or 40x magnification. 3D printed cell-laden grids were incubated in media for 5 days, removed, and fresh 3 ml media supplemented with either 1 mM theophylline or a 1% DMSO vehicle control was added to the gel. After 24 hours, the patterns were removed from the solutions and a subsection of each gel was cut for imaging. Brightfield, chlorophyll autofluorescence, and YFP fluorescence images were captured for each sample containing the different YFP strains. A TRITC channel was used to monitor in vivo chlorophyll autofluorescence from cyanobacteria.

### YFP monitoring by fluorescence measurements from surrounding medium

To assess the presence of YFP in the surrounding medium, the 24-hour theophylline/DMSO supplemented samples (as prepared above) were analyzed using fluorescence measurements. The surrounding media of the hydrogel was collected and centrifuged at 4500 x g for 5 min and the emission intensity of YFP from the supernatant was measured in a 96-well plate with a Tecan Infinite M200 plate reader (TECAN). The excitation and emission wavelengths were set for YFP to excitation 490/9 nm and emission 535/20 nm. Measurements were taken from hydrogels containing 3 independent clones of each YFP strain along with the WT cell-loaded sample.

### SDS-PAGE of CotA

Cultures of WT and Laccase^+^-riboF-Lysis^+^ were grown in liquid culture under 75 μmol photons m^-2^ s^-1^ with continuous shaking at 30 °C to an OD_750_ of 0.5. 10 mL of cultures were pelleted by centrifugation at 4500 x g and aspirated. The pellets were then resuspended in 1 mL of cold lysis buffer (50 mM Tris-HCl pH 8.0, 150 mM NaCl) supplemented with 1 mM PMSF and Complete Mini Protease Inhibitor (Roche). The samples were then homogenized in a BeadBlaster 24R (Benchmark Scientific) at 4 °C. The crude protein was then clarified by centrifugation at 4 °C for 20 min at 18,000 x g, after which the crude protein concentration was determined using a Pierce Coomassie Plus Protein Assay (Bradford), with bovine serum albumin used as a standard. After boiling the sample with 2x Laemmli Sample Buffer, 30 μg of sample was loaded into a Mini-PROTEAN® TGX Precast Gel (Bio Rad) and run at 150V. The gel was then incubated in InstantBlue Coomassie Protein Stain (Abcam) overnight prior to imaging.

### ABTS activity assays

Laccase activity of either the hydrogel or the surrounding solution was measured using ABTS (Thermo Fisher Scientific). For assays involving the induction of RiboF-Laccase^+^, an approximately 83 mm^3^ subsection of a hydrogel (two interconnected hollow squares) was added to 1 mL of BG-11 in a 2 mL Eppendorf tube and supplemented with either 1 mM theophylline (Sigma-Aldrich; 100 mM stock dissolved in 100% DMSO) or 1% DMSO as a control and allowed to incubate for 3 days under 50 μmol photons m^-2^ s^-1^ prior to conducting the ABTS assays. The hydrogel sample was then transferred to a 1 mL reaction buffer mixture of 100 mM sodium acetate buffer, 3.16 μM CuSO_4_, pH 5.0, and 2 mM ABTS in a 1.5 ml conical microtube and incubated in darkness at 28 °C. The oxidation of ABTS was determined by measuring the absorbance of the reaction mixture at 420 nm^53^ after 24 hours. A blank reading at 420 nm of the reaction mixture without a hydrogel was subtracted from all samples.

For ABTS assays within the hydrogels with Laccase^+^-riboF-Lysis^+^ cells, an approximately 83 mm^3^ subsection of the hydrogel was submerged in 1 ml of reaction buffer mixture (as mentioned above) and incubated for 1 hour. The hydrogel ELMs were then removed from the solution and placed on sterile agar LB plates and incubated for 4 days in a dark environment at 28 °C. Photographs were taken at regular time intervals. For laccase activity in the surrounding solution, similar subsections of the hydrogels were submerged in 1 ml of the reaction buffer (as mentioned above) containing 2 mM ABTS and incubated for 4 days in a darkness at 28 °C. Absorbance of the reaction mixture was measured at 420 nm at regular time intervals, relative to the absorbance at 0 h. A blank reading at 420 nm consisting of the reaction mixture without a hydrogel was subtracted from all samples. Additionally, for the Laccase^+^-riboF-Lysis^+^ hydrogel set, supernatant laccase activity was measured. 1 mL of the medium surrounding a 5-day grown hydrogel ELM was centrifuged for 5 minutes at 4500 x g. 500 μL of the supernatant was combined with 460 μL 200 mM sodium acetate buffer, pH 5.0, and 40 μL of a 50 mM stock of ABTS for a final concentration of 100 mM sodium acetate buffer, pH 5.0, and 2 mM ABTS. The samples were incubated in darkness at 28 °C, and oxidation of ABTS was measured at regular time intervals, relative to the absorbance at 0 h. A blank reading at 420 nm consisting of the BG-11 in the reaction mixture was subtracted from all supernatant samples. For all the assay measurements, 200 μL of samples were taken in a 96-well plate and measured with a Tecan Infinite M200 plate reader (TECAN).

### Indigo carmine decolorization assays

The decolorization of indigo carmine by hydrogels was determined by suspending an 83 mm^3^ subsection of a hydrogel (two interconnected hollow squares) in a 500 μL solution of BG-11 media with 0.1 mM indigo carmine (Sigma-Aldrich). Indigo carmine concentrations were determined spectrophotometrically by generating a standard curve for absorbance at 612 nm for 200 μL samples in a 96-well plate with a Tecan Infinite M200 plate reader. The indigo carmine hydrogel mixture was incubated in darkness, and the absorbance of 200 μL samples was measured at 612 nm at time 0, and intermittently till 10 days of incubation at 28 °C. The percentage of dye decolorization after 10 days was determined relative to absorbance at 612 nm at time 0 for a given sample.

### Inducible cell death by phage-gene lysis within 3D printed hydrogel

Small grid_B hydrogel patterns with strains Laccase^-^-riboF-Lysis^-^ and Laccase^+^-riboF-Lysis^+^ were printed and incubated for 5 days of growth. After 5 days, the constructs were transferred to flasks containing 5 ml of either BG11 medium, 1 mM theophylline (prepared from a 100 mM stock dissolved in 100% DMSO), or 1% DMSO as a control. The flasks were kept under 75 μmol photons m^-2^ s^-1^ with continuous shaking at 30 °C. Images of the experimental samples were captured on day 0 and day 12.

### Inducible cell death by phage-gene lysis in liquid culture

Strains Laccase^-^-riboF-Lysis^-^ and Laccase^+^-riboF-Lysis^+^ were grown in liquid culture under 75 μmol photons m^-2^ s^-1^ with continuous shaking at 30 °C to an initial OD_750_ of 0.2 - 0.3, prior to being transferred to six-well plates in 8 ml aliquots. Samples were then induced with either 1 mM theophylline (prepared from a 100 mM stock dissolved in 100% DMSO) or 1% DMSO as a control. The OD_750_ of the cultures were measured daily for four days following induction.

### Construction of plasmids and engineered strains

To create plasmid pAM5823, pAM4950 was digested with SwaI (NEB) to release the *ccdB* toxic gene, and the linearized plasmid was re-ligated with a Quick Ligase Kit (NEB) following manufacturer instructions. Plasmid pAM5825 were created by digesting pAM4950 with SwaI and performing a Gibson Assembly (NEB) between the linearized product and a *cotA* gBlocks Gene Fragment synthesized by Integrated DNA Technologies (IDT). For Gibson Assembly, the *cotA* gBlocks gene fragment contained 20 base pair ends of homology with the linearized pAM4950 product and a *con*II promoter and ribosomal binding site sequence derived from pAM4909 upstream of the *cotA* gene. The *cotA* sequence was derived from a reverse-translation of the protein sequence of CotA from *Bacillus subtilis* (accession number WP_003243170.1) and was codon optimized for *S. elongatus*. pAM5826 was created by digesting pAM5051 with EcoRI (NEB), and performing a Gibson Assembly between the linearized product and the PCR product of pAM5825 with primer pair cotA_5051-F/cotA_5051-R. pAM5829 was created by digesting pAM4940 with SwaI and performing a Gibson Assembly of the linearized product and the PCR product of *S. elongatus* gDNA and the primer pair 0766_riboF_pAM4940-F/0766_riboF_pAM4940-R. The sequence of all plasmids constructed in this study were verified by Sanger sequencing. Complete lists of all plasmids and primers used in this study are included in Tables S2 and S3, respectively. Plasmids pAM4909, pAM5027, and pAM5057 were transferred into *S. elongatus* by biparental conjugation^54^. Transformation of *S. elongatus* with all other plasmids was achieved using standard protocols^55^. All *S. elongatus* transformations were performed using WT strain AMC06. Genotyping of *S. elongatus* was performed using colony PCR with Q5 DNA Polymerase (NEB), using primer pair NS1_Screen-F/NS1_Screen-R for transformations in neutral site 1 (NS1), and NS2_Screen-F/NS2_Screen-R for transformation in neutral site 2 (NS2). A complete list of strains used in this study is included in Table S4.

### Statistical Analysis

Data are presented as the mean ± standard deviation unless noted otherwise. The definition of significance was set a priori to p < 0.05. Within figures, asterisks represent: *, P < 0.05; **, P < 0.01; ***, P < 0.001; ****, P < 0.0001.

## Supplemental Information

Supporting figures and tables are listed in the Supplementary information.

## Acknowledgments

We thank David Wirth for assistance with the creation of the 3-way syringe mixer, Jack Reddan for assistance with the construction of pAM5825, Marie Adomako for the construction of pAM5610, and Bryan Bishé and Arnaud Taton for assistance with the conjugation of strains YFP^+^, YFP^-^, and Ribo-YFP. This work was primarily sponsored by the UC San Diego Materials Research Science and Engineering Center (UCSD MRSEC), supported by the National Science Foundation (Grant DMR-2011924). The authors acknowledge the use of facilities and instrumentation supported by the National Science Foundation through the UCSD MRSEC DMR-2011924, the Department of NanoEngineering Materials Research Center (NE-MRC), the UCSD School of Medicine Microscopy core facility funded by NINDS Grant P30 NS047101, and the Gordon and Betty Moore Foundation Aquatic Symbiosis Model Systems (Grant 9325 to D.W. and S.C.). The use of microscopy core equipment was supported by Jennifer Santini and Marcella Erb.

