## Supplemental Information for "Phenotypically Complex Living Materials Containing Engineered Cyanobacteria"

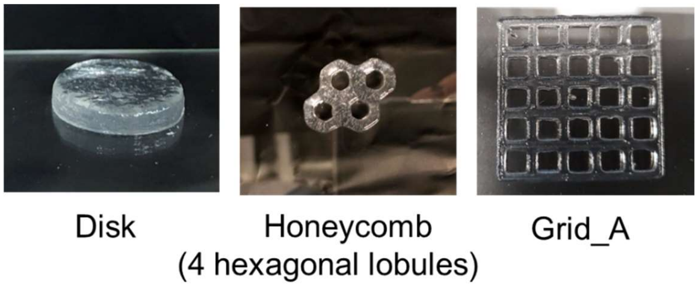


**Figure S1:** Different patterns of 3D printed alginate hydrogel.

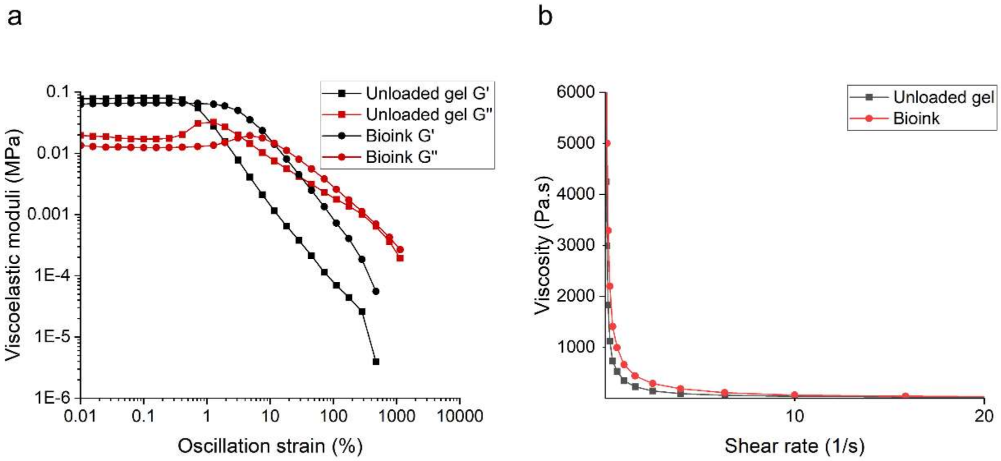


**Figure S2.** Rheological characterization of the unloaded gel and bioink after five days of incubation. a) Viscoelastic moduli vs. oscillatory strain at ω = 10 rad/s and 25 °C. Storage modulus is denoted by G' and loss modulus by G''. b) Viscosity vs. shear rate of hydrogels at 25 °C.

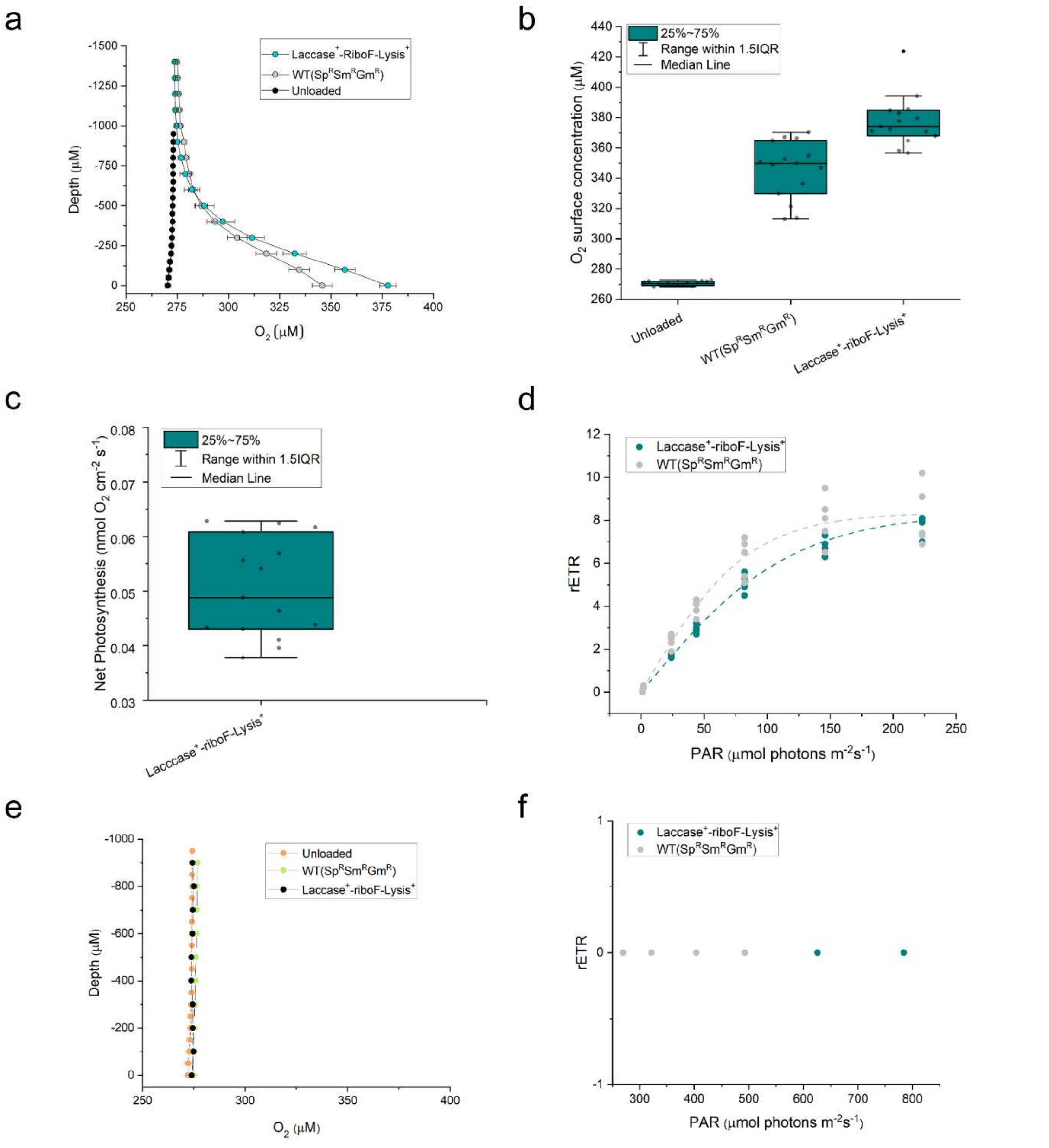


**Figure S3**. $O_2$ microenvironment and photosynthetic activity of living hydrogels. a) $O_2$ microsensor profiles were performed from the hydrogel surface (depth = 0 μm) into the overlying water column for an unloaded hydrogel control, a hydrogel containing the WT($Sp^RSm^RGm^R$) (labeled Laccase$^-$-riboF-Lysis$^-$) strain and a hydrogel containing the Laccase$^+$-riboF-Lysis$^+$ strain. Data are means ± SEM (n=15 measurements from the 3 independent hydrogels). b) Box plot of $O_2$ concentrations measured at the surface of the hydrogel. c) Box plot of net photosynthesis at an incident downwelling irradiance of 80 μmol photon $m^{-2}$ $s^{-1}$ for Laccase$^+$-riboF-Lysis$^+$ cells encapsulated in the hydrogel. d) Relative electron transport (rETR) vs. photosynthetically active radiation (PAR; 400-700 nm) measured as rapid light curves for 5 replicate spots for each printed hydrogel of the mutant WT($Sp^RSm^RGm^R$) and Laccase$^+$-riboF-Lysis$^+$ strains. Dotted lines indicate non-linear curve fits according to the Platt photosynthesis model (see methods) ($r^2 > 0.95$). e) $O_2$ microsensor measurements during darkness. $O_2$ microsensor profiles were performed for an unloaded hydrogel control, a hydrogel containing the WT($Sp^RSm^RGm^R$) strain, and a hydrogel containing the Laccase$^+$-riboF-Lysis$^+$ strain. f) rETR vs. PAR data above 250 μmol photons $m^{-2}$ $s^{-1}$ showed full photoinhibition.

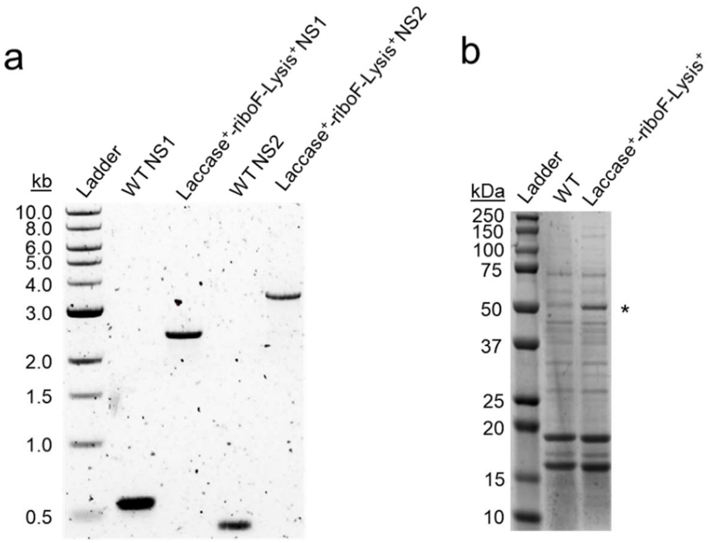


**Figure S4.** Characterization of the strain Laccase$^+$-riboF-Lysis$^+$ a) Agarose gel showing the genotypic characterization of the Laccase$^+$-riboF-Lysis$^+$ strain. Lane 1, standard 1-kb ladder (NEB); lane 2, PCR amplification of WT gDNA with primers surrounding neutral site 1; lane 3, PCR amplification of Laccase$^+$-riboF-Lysis$^+$ gDNA with primers surrounding neutral site 1; lane 4, PCR amplification of WT gDNA with primers surrounding neutral site 2; lane 5, PCR products of Laccase$^+$-riboF-Lysis$^+$ gDNA with primers surrounding neutral site 2. b) SDS-PAGE of protein extracted from cultures of WT and Laccase$^+$-riboF-Lysis$^+$ strains grown at 30 °C on an orbital shaker under 70 μmol photon $m^{-2}$ $s^{-1}$, with the band corresponding to the predicted size of CotA indicated by an *.

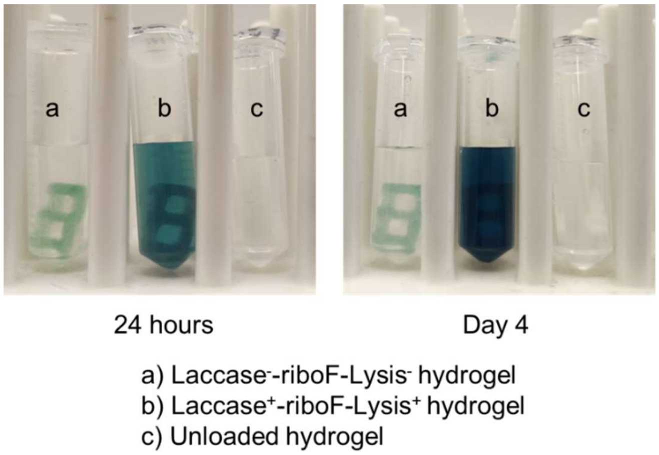


**Figure S5.** Images of 3D printed patterns incubated in ABTS reaction buffer.

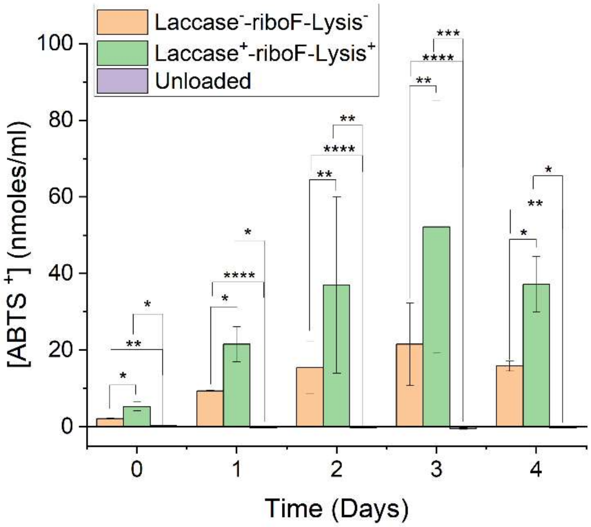


**Figure S6.** Laccase activity of the supernatant of media surrounding hydrogels. Time-course of the oxidation of ABTS in reaction buffer with the addition of supernatant surrounding either the Laccase$^{-}$-riboF-Lysis$^{-}$, Laccase$^{+}$-riboF-Lysis$^{+}$ strain, or an unloaded hydrogel.

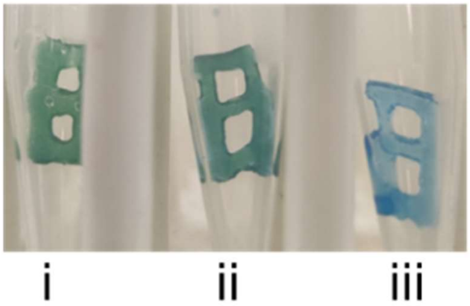


**Figure S7.** Hydrogels separated from solutions after 10 days of incubation. Roman numerals indicate the addition of i) a hydrogel printed with Laccase$^{+}$-riboF-Lysis$^{+}$, ii) a hydrogel printed with Laccase$^{-}$-riboF-Lysis$^{-}$, iii) unloaded hydrogel.

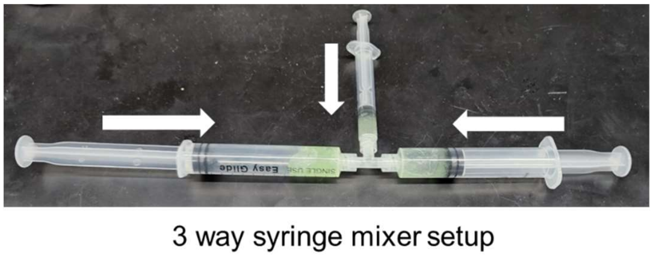


**Figure S8.** In-house fabricated 3-way syringe mixer for bioink preparation.

**Table S1:** Dimension and layer information of 3D printed gel patterns

| **Pattern Type** | **Dimensions** | **Number of Layers** |
|---|---|---|
| Disk | Radius: 10 mm<br>Z height: 5 mm | 13 |
| Honeycomb (4 hexagonal lobules) | Inner length: 4 mm<br>Outer length: 5 mm<br>Z height: 5 mm | 17 |
| Honeycomb (7 hexagonal lobules) | Inner length: 4 mm<br>Outer length: 5 mm<br>Z height: 5 mm | 13 |
| Grid_A | Length X and Y: 29 mm<br>Z height: 1.5 mm | 5 |
| Grid_B | Length X and Y: 12.5 mm<br>Z height: 1.5 mm | 5 |

**Table S2.** Plasmids used in this study.

| Plasmid Name | Description | Source |
|---|---|---|
| pAM4909 | RSF1010 broad host-range plasmid with *aadA* cassette and constitutive expression of *myc-yfp* reporter driven by a PconII promoter. | Taton *et al.* 2014 |
| pAM4940 | Contains recombination arms for NS1 integration, an *aacC1* cassette, and a *ccdB* gene surrounded by SwaI restriction sites. | Taton *et al.* 2014 |
| pAM4950 | Contains recombination arms for NS2 integration, an *aadA* cassette, and a *ccdB* gene surrounded by SwaI restriction sites. | Taton *et al.* 2014 |
| pAM5027 | RSF1010 broad host-range plasmid with *aadA* cassette. | Ma *et al.* 2014 |
| pAM5051 | Plasmid containing recombination arms for NS2 integration and an *aadA* cassette, with a PconII promoter driving the expression of riboswitch-F followed by an EcoRI restriction site. | Ma *et al.* 2014 |
| pAM5057 | RSF1010 broad host-range plasmid with *aadA* cassette and riboswitch-F::*myc-yfp* reporter expression driven by a PconII promoter. | Ma *et al.* 2014 |
| pAM5329 | Plasmid containing recombination arms for NS1 integration and an *aadA* cassette. | Taton *et al.* 2017 |
| pAM5610 | Plasmid containing recombination arms for NS1 integration and an *aacC1* cassette. | This study |
| pAM5823 | Plasmid containing recombination arms for NS2 integration and an *aacC1* cassette. | This study |
| pAM5825 | Constitutive expression of *cotA*, driven by a PconII promoter, with recombination arms NS2 integration and an *aadA* cassette. | This study |
| pAM5826 | Plasmid containing a PconII promoter driving the expression of riboswitch-F::*cotA*, with recombination arms for NS2 integration and an *aadA* cassette. | This study |
| pAM5829 | Plasmid containing a PconII promoter driving the expression of riboswitch-F::*Synpcc7942_0766* driven by a PconII promoter, with recombination arms for NS1 integration, and an *aacC1* cassette | This study |

**Table S3** Primers used in this study.

| **Primer Name** | **Sequence** |
|---|---|
| cotA_5051-F | CTGCTAAGGAGGCAACAAGATGACGCTGGAAAAGTTCGTCGA |
| cotA_5051-R | CGAGGTCGAGACGGCTCATGTTATTTGTGGGGATCGGTGA |
| 0766_riboF_pAM4940-F | GGCCAATAACCCAGGGATTTTTGACAATTAATCATCGGCTCG |
| 0766_riboF_pAM4940-R | CTGCCGGGGAGCTCCTTCATTTCATGTCAGGGTTTCAGGCAG |
| NS1_Screen-F | ACATCGCTATCTCTTAGGACTTCG |
| NS1_Screen-R | GGCCGAAAATGACAAGATCAC |
| NS2_Screen-F | CTCCAGTAAAGTCTTCGCCCGTAAC |
| NS2_Screen-R | TTGGTGCTGTTCAGTCTGGATGC |

**Table S4.** Strains used in this study.

| **Name** | **Description** | **Plasmids Used** | **AMC #** | **Source** |
|---|---|---|---|---|
| WT | Wild-type *S. elongatus* (AMC06) | | AMC06 | |
| $YFP^{+}$ | Harbors RFS1010 plasmid for constitutive expression of *myc-yfp* | pAM4909 | AMC2771 | This study |
| $YFP^{-}$ | Harbors RFS1010 plasmid. Negative control for $YFP^{+}$ and RiboF-YFP. | pAM5027 | AMC2772 | This study |
| RiboF-$YFP^{+}$ | Harbors RSF1010 plasmid with *myc-yfp* expression regulated by riboswitch-F. | pAM5057 | AMC2773 | This study |
| RiboF-$Laccase^{+}$ | Contains PconII driving cotA expression regulated by riboswitch-F in NS1. | pAM5826 | AMC2738 | This study |
| RiboF-$Laccase^{-}$ | Negative control strain for RiboF-$Laccase^{+}$. | pAM5329 | AMC2734 | This study |
| $Laccase^{+}$-riboF-$Lysis^{+}$ | Contains PconII driving constitutive expression of cotA in NS2, and PconII driving expression of *SynPCC7942_0766* regulated by riboswitch-F in NS1. | pAM5825, pAM5829 | AMC2740 | This study |
| $Laccase^{-}$-riboF-$Lysis^{-}$<br>or<br>WT ($Sp^{R}Sm^{R}Gm^{R}$) | Negative control strain for Laccase+-riboF-Lysis+. | pAM5610, pAM5823 | AMC2739 | This study |

### Supplementary Text

*Photosynthetic activity of strain $Laccase^{+}$-riboF-$Lysis^{+}$*

Similar to the control strain, $O_2$ microsensors revealed a hyperoxic microenvironment on the surface of the living hydrogels that carried the embedded $Laccase^{+}$-riboF-$Lysis^{+}$ strain (**Figure S3a & b**). $O_2$ concentrations were slightly higher for the $Laccase^{+}$-riboF-$Lysis^{+}$ hydrogel (mean=378 ± 4.2 µM SE); (Tukey post hoc p = <0.01) compared to the WT($Sp^{R}Sm^{R}Gm^{R}$) hydrogel at an incident irradiance of 80 µmol photons $m^{-2}$ $s^{-1}$. Net photosynthesis in the $Laccase^{+}$-riboF-$Lysis^{+}$ strain (**Figure S3c)** was approximately 1.6-fold

(0.05 nmol $O_2$ $cm^{-2}$ $s^{-1}$) that of the WT($Sp^RSm^RGm^R$) strain (0.031 nmol $O_2$ $cm^{-2}$ $s^{-1}$, Tukey post hoc, $p < 0.01$). During darkness hydrogels that encapsulate the Laccase$^+$-riboF-Lysis$^+$ strain showed limited respiratory activity and $O_2$ concentrations were close to ambient seawater values as also observed for the control strain (**Figure S3e**). Like the WT($Sp^RSm^RGm^R$) *S. elongatus* strain, measurements of variable chlorophyll a fluorimetry for the Laccase$^+$-riboF-Lysis$^+$ strain was saturated between 200 – 250 µmol photons $m^{-2}$ and indicative of low light adaptation with an irradiance at onset of saturation of about 120 µmol photons $m^{-2}$ $s^{-1}$ for the laccase mutant strain (**Figure S3d**).
